# microRNA Expression Levels Change in Neonatal Patients During and After Exposure to Cardiopulmonary Bypass

**DOI:** 10.1101/2021.08.13.454953

**Authors:** Lance Hsieh, Lan Tu, Alison Paquette, Nataliya Kibiryeva, Jennifer Marshall, Douglas Bittel, James O’Brien, Kasey Vickers, Peter Pastuszko, Vishal Nigam

## Abstract

**Objectives:** The systemic inflammation that occurs after exposure to cardiopulmonary bypass (CPB), which is especially severe in neonatal patients, is associated with poorer outcomes and is not well understood. In order to gain deeper insight into how exposure to bypass activates inflammatory responses in circulating leukocytes, we studied changes in microRNA (miRNA) expression during and after exposure to bypass. miRNAs are small non-coding RNAs that have important roles in modulating protein levels and function of cells.

**Methods:** We performed miRNA-Sequencing on leukocytes isolated from neonatal cardiopulmonary bypass patients (N=5) at 7 timepoints during the process of CPB, including prior to the initiation of bypass, during bypass, and at three time points during the first 24 hours after weaning from bypass. We identified significant differentially expressed miRNAs using generalized linear regression models, and miRNAs were defined as statistically significant using an FDR adjusted p <0.05. We identified gene targets of these miRNAs using the Targetscan database, and identified significantly enriched biological pathways for these gene targets.

**Results:** We identified 54 miRNAs with differential expression during and after CPB. These miRNAs clustered into 3 groups, including miRNAs that were increased during and after CPB (3 miRNAs), miRNAs that decreased during and after CPB (10 miRNAs), and miRNAs that decreased during CPB but then increased 8-24 hours after CPB. 38.9% of the target genes of these miRNAs were significantly differentially expressed in our previous study. miRNAs with altered expression levels are predicted to significantly modulate pathways related to inflammation and signal transduction.

**Conclusions:** The unbiased profiling of the miRNA changes that occur in the circulating leukocytes of bypass patients provides deeper insight into the mechanisms that underpin the systemic inflammatory response that occurs in patients after exposure to cardiopulmonary bypass. These data will help the development of novel treatments and biomarkers for bypass associated inflammation.

## INTRODUCTION

Surgical palliation/repair of congenital heart defects is associated with significant morbidity, mortality, and cost. Neonatal patients are particularly at risk, having a 10% mortality and a 30% complication rate.^1, 2^ The vast majority of pediatric open-heart surgeries require the patient to be supported by cardiopulmonary bypass (CPB) to give the surgeon a bloodless field to operate in, while also minimizing ischemic damage to the body. However, CPB patients experience significant post-CPB inflammation—which includes increased cytokine levels, inflammatory cell infiltration, vascular leak, and multi-organ dysfunction. In neonates recovering from complicated cardiac surgeries, increased cytokine levels are associated with high mortality and extended intensive care stays.^3^ However, the molecular mechanisms that underlie this inflammatory response are not fully understood.

In an effort to understand how CPB instigates inflammation, we have quantified functional changes in circulating leukocytes using unbiased and comprehensive ‘omics approaches. Recently, we have measured global gene expression changes that occur in neonatal patients during and after CPB exposure using RNA sequencing^4^. In this report, we examine the microRNA (miRNA) changes that occur in neonatal CPB patients. miRNAs are small non-coding RNAs that bind to specific regions of mRNAs based on underlying sequence, which are then known as “target genes”^5^. miRNAs modulate the protein levels of their target genes by either attenuating mRNA translation and/or promoting degradation of the target mRNAs^6^. Through disruption of these target genes, miRNAs play a vital role in numerous cellular and biological processes, including inflammatory signaling, and have been demonstrated to be differentially expressed in many human diseases, such as cardiovascular diseases^7^.

There have been a lot of efforts and interest in finding miRNA biomarkers for post CPB complications. For example, miRNAs have been studied as biomarkers of myocardial injury^8, 9^. Currently, a limited amount is known about the miRNA changes that in leukocytes in response to CPB. This knowledge gap is significant since leukocytes are crucial regulators of inflammation. So, understanding global changes of miRNA in leukocytes is a critical step to better understand the mechanisms of CPB induced inflammation, and how these miRNAs may regulate this inflammation.

We performed a comprehensive unbiased profiling of miRNA expression from pediatric patient leukocytes. Blood samples were collected pre, during, and post bypass surgery at 7 different timepoints from 5 infants with CHDs undergoing CPB surgery. Leukocytes were isolated from whole blood from each set of patient samples miRNA was quantified using RNA sequencing. This study aimed to characterize CPB affected leukocyte miRNA profiles and to holistically analyze how differential expression of these miRNAs may impact target gene expression and corresponding changes in biological pathways.

## METHODS

### Human blood samples

Pediatric patients less than 1 month old with different congenital heart defects requiring repair utilizing CPB were enrolled in our study at Mercy’s Children Hospital (Kansas City, Missouri, USA). Written informed consent was received from participants’ parents or legal guardians prior to inclusion in the study. A standard CPB protocol was utilized in all of the patients. Approach in all cases was via median sternotomy. Aortic cannulation was used for the arterial access, with either single venous cannula being placed in the right atrium or bicaval cannulation, depending on the type of the defect requiring repair. The CPB circuit was blood primed, and standard additives included Solu Medrol, sodium bicarbonate, cefazolin, and tranexamic acid. After initiation of CPB, the patients were cooled to 18°C–30°C. Mean perfusion pressure was maintained at 25–35 mmHg, and the minimum hematocrit was kept at 24%. Upon completion of repair, the patients were rewarmed and weaned from CPB. Modified ultrafiltration (MUF) was performed in all of the patients after weaning from CPB. Blood samples of 2.5 mL were collected from an indwelling patient line or from the CPB pump, dependent upon the time of the blood draw (for blood draws performed while the patient was on CPB, these samples were pulled from the CPB pump). These samples were collected in EDTA tubes at 7 time points: before CPB (CPB-0h), 1 hour into CPB (CPB-1h), end of CPB (CPB-end), end of MUF (MUFend), and 1 hour (MUF-1h), 8 hours (MUF-8h), 24 hours (MUF-24h) during postoperative recovery. Blood was centrifuged at 1,000*g* for 20 minutes at room temperature, and plasma was removed. RBC were lysed in RNase-free RBC lysis solution (PerfectPure RNA blood kit, 5Prime) for 5 minutes at room temperature. The tubes were centrifuged at 2000*g* for 5 minutes at room temperature. The pellets of nucleated cells were lysed in the cell lysis buffer and immediately stored at −80°C till RNA extraction. The design and time course of the study are illustrated in **Figure 1A**. Patient demographics are in **Table 1.**

**Figure 1:**
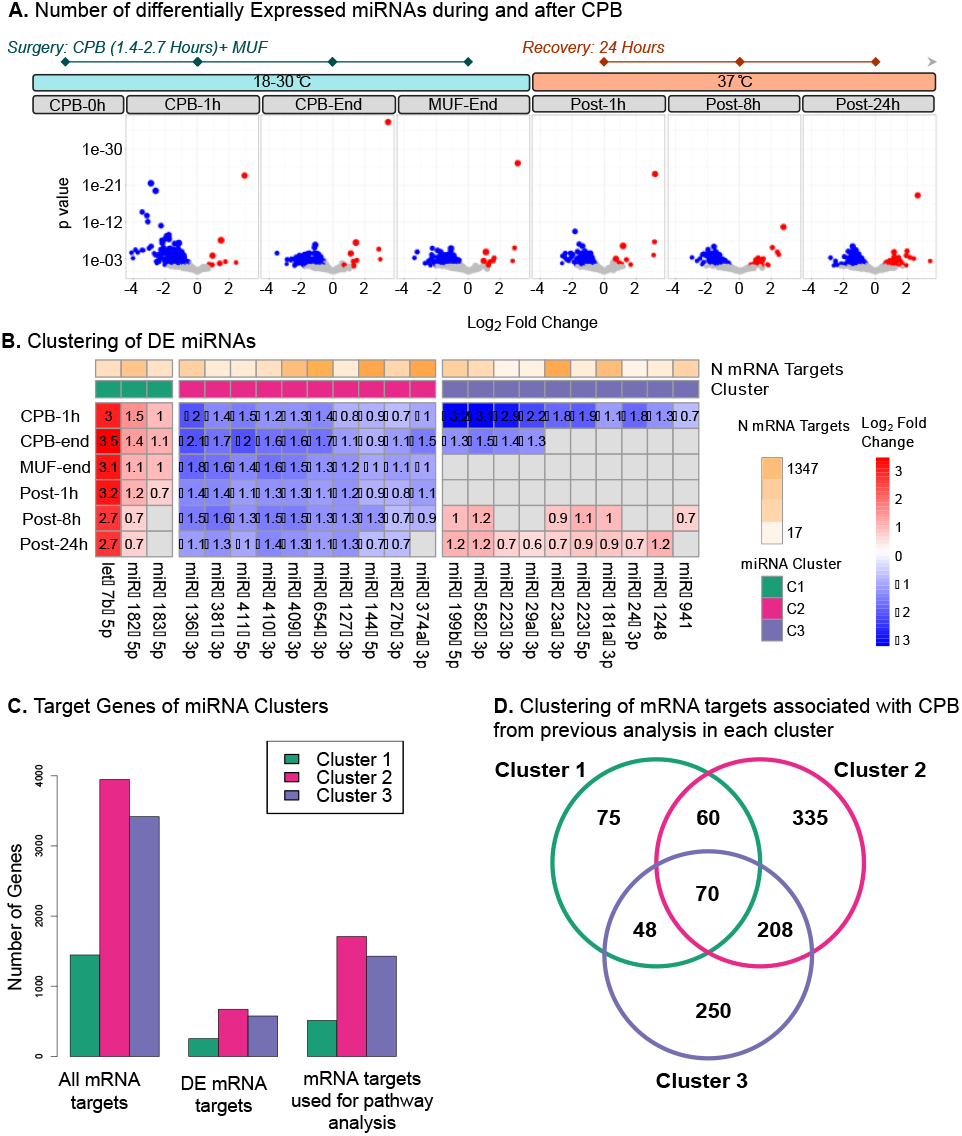
Changes in miRNA expression during and after CPB. **(A)** The design and time course of the study. Neonates with different congenital heart diseases underwent CPB surgery that lasted 1.4-2.7 hrs on average, followed by a short duration of modified ultrafiltration (MUF). Body temperature was cooled down to 18-30°C during surgery and quickly rewarmed to 37°C after MUF. Blood samples were collected at 7 time points; miRNA from isolated nucleated cells were submitted for sequencing (n=5 patients). The number of miRNAs that were significantly downregulated or upregulated at different time points compared to the expression level before surgery (CPB-0h). Significance was defined as FDR adjusted p-value <0.05. **(B)** 54 miRNAs were divided into 3 clusters: cluster 1 with miRNAs upregulated in both phases, cluster 2 with miRNAs downregulated in both phases, and cluster 3 with miRNAs downregulated in the surgery phase but upregulated in the recovery phase. The heatmap shows the log_2_ fold changes miRNAs in each cluster compared to before CPB. **(C)** Bar plot showing the total number of mRNA targets within each cluster. A subset of the targets was previously associated with CPB.^4^ Only the top 200 targets for each miRNA was used for KEGG Pathway analysis (Figure 2). the number of mRNA targets for each individual miRNA are presented in supplemental table 2. (D) Overlapping and distinct target genes within each cluster that were previously associated with CPB.^4^

**Table 1 –.** Patient Demographics.

| **I**<br>**D** | **Age**<br>**(days)** | **Weight**<br>**(kg)** | **Sex** | **Diagnosis** | **Lowest temp**<br>**(°C)** | **Surgery**<br>**time (min)** | **Bypass**<br>**time (min)** |
| --- | --- | --- | --- | --- | --- | --- | --- |
| 1 | 5 | 3.3 | M | TGA, IVS, ASD | 27.4 | 270 | 161 |
| 2 | 5 | 3.3 | M | HLHS, PDA, ASD | 18.2 | 234 | 158 |
| 3 | 19 | 2.7 | F | TAPVC | 18.0 | 163 | 86 |
| 4 | 12 | 3.4 | F | TAPVC | 21.4 | 173 | 99 |
| 5 | 4 | 3.9 | M | HLHS, PDA | 17.6 | 295 | 161 |
| TGA – Transposition of the great artery, IVS – Intact ventricular septum, ASD – Atrial septal defect, HLHS – Hypoplastic left heart syndrome |  |  |  |  |  |  |  |

### miRNA Sequencing

Total RNA from the nucleated cells was extracted using either the MirVana RNA extraction kit (Invitrogen). The concentration and quality of total RNA for each sample were quantified and evaluated by spectrophotometry (Epoch; Thermo Scientific) and automated chip electrophoresis analyses (Experion, BioRad) respectively. Each sample met the quality standards of a 260/280 ratio > 2.0 and a RNA integrity number (RIN) > 9. Illumina TruSeq RNA libraries were prepared from 1 μg total RNA and sequenced to generate 50 base pair single-end reads using the the HiSeq 1500 (Illumina, USA) at the Children’s Mercy OMICs Research Core Lab (Kansas City, MO).

### Bioinformatics

Small RNA sequencing datasets were analyzed using the TIGER pipeline.^10^ Briefly, raw reads were preprocessed for quality control using FastQC (www.bioinformatics.babraham.ac.uk/projects/fastqc). Cutadapt (v2.9) was used to trim 3’ adapters and reads <16 nts in length were removed.^11^ Preprocessed sRNA reads were collapsed into non-redundant identical files and aligned to the human genome using bowtie (v1.2.3) allowing 1 mismatch.^12^ For miRNAs, non-templated additions were clipped from the 3’ terminal end and sRNA reads overlapping mature miRNA coordinates were counted allowing for offset positions −2, −1, 0, +1, +2. Results were reported as raw read count and reads per million total reads (RPM). Differential expression analyses were performed by DEseq2 (v1.24.0).^13^ We performed adjustment for multiple comparisons using the Benjamini-Hotchberg approach^14^. miRNAs were considered statistically significant at a false discovery rate (FDR) adjusted *p* <0.05. miRNAs were considered to be significantly altered during either surgery or recovery if there were significant differences for at least one timepoint compared to baseline within each respective stage of CPB (surgery or recovery). A cutoff of 25 reads per million (RPM) was used to filter out any miRNAs with low expression. Differentially expressed miRNAs were manually clustered based on the directionality of the log fold change, and visualized using the R package “pheatmap” (Version 1.0.12)(https://www.rdocumentation.org/pack-ages/pheatmap/versions/1.0.12/topics/pheatmap-package).

### Identification of Putative miRNA targets

Putative mRNA targets for miRNAs within each cluster were detected using the quantitative model TargetScan (V. 7.2, targetscan.org), which characterizes canonical targeting of miRNAs based on 14 features, and has the best predictive performance compared to comparable tool.^15^ For clusters which contained >10 miRNAs, we included the top 10 miRNAs based on log FC change. We removed all mRNA targets that were not present in the paired RNA sequencing data (N=12,709 genes). We performed additional filtering using the absolute value of the “context++” score, which is a metric of miRNA target prediction accuracy used by TargetScan.^15^ For each miRNA we removed all putative targets with a context++ score of 0. We also eliminated all mRNAs that were not present in our ancillary mRNA analysis.^4^ This filtering ensured that only the highest quality relationships were validated.

### Pathway Enrichment Analysis of miRNA Target Genes

We performed pathway enrichment analysis of the genes regulated by each cluster of miRNAs using the EnrichR R package (Version 2.1)^16^.For this analysis, we included only the top 200 genes (based on context++ score) for each miRNA. Through Enrichr, we analyzed KEGG biological pathways (excluding KEGG “Human Disease” pathways). Pathways were considered statistically significant with an FDR adjusted p <0.05. miRNA-gene-pathway networks were visualized using cytoscape (Version 3.7.2) ^17^. All data was analyzed and visualized using R (Version 4.0.3).

### Cytokine-Receptor Networks

A subnetwork of Receptor-Ligand networks were generated for target genes using the Fantom5 database ^18^; which contained 708 ligands and 691 receptors. We performed a sub analysis of predefined cytokines, chemokines and integrin/ adhesion molecules from within this receptor ligand network, based on the class of molecules known to be mediators of inflammation.

## RESULTS

### Micro RNA sequencing analysis of total patient nucleated cells throughout CPB

To characterize the miRNA expression profiles of patient blood affected by CPB, miRNA sequencing analysis was performed on RNA isolated miRNA from total nucleated cells of 5 neonate patients undergoing CPB surgery (**Table 1**). Neonates with varying CHDs (Table 1) underwent CPB surgery procedure (bypass time range 86-161 minutes, mean bypass time 133 Minutes, SD 37.33 minutes), outlined in **Figure 1A**, with blood samples collected at 7 different time points prior to, during, and after CPB. miRNAs contributed to >60% of the small non-coding RNAs present in the blood collected during these times (**Supplemental Figure S1A**). Detection of unique miRNAs were characterized as normalized expression in reads per million (RPM) averaged over all 5 patient samples, within each CPB timepoint. The 4 most abundant miRNAs across all timepoints were hsa-mir-486-5p, hsa-mir-92a-3p, hsa-mir-451a, and hsa-mir-16-5p, (**Supplemental Figure S1B**).

### Identification of differentially expressed miRNAs in different stages of CPB

To obtain a better understanding of how CPB affects miRNA expression, we compared the expression of miRNAs at surgery and recovery timepoints to the pre CPB timepoint (CPB-0h). Differentially expressed miRNAs were identified through generalized linear regression models within DESEQ2.^12^ Within each timepoint, we observed that there were more differentially expressed miRNAs that were significantly downregulated compared to baseline, than miRNAs that were upregulated compared to baseline (**Figure 1A**). 42 of these miRNAs had significantly altered expression at one timepoint during surgery compared to baseline, 54 miRNAs had significantly altered expression both during bypass and after bypass, and 13 miRNAs had altered expression compared to pre-CPB only in the recovery from CPB phase, using a cutoff for significance of FDR adjusted p <0.05 and a RPM of >25.

In order to determine the effect of CPB on miRNA expression, the 54 miRNAs had significant modulation of expression at least one timepoint in both surgery and recovery were clustered into 3 groups based on their expression patterns throughout the course of CPB and recovery (**Figure 1B**). One cluster (Cluster 2) had a total of 41 miRNAs significantly decreased using our clustering threshold, and so we only included the top 10 miRNAs based on fold change. Cluster 1 (green) represents the miRNAs that remained significantly up regulated in both CPB and recovery phases (Fig 1B), cluster 2 (pink) represents the miRNAs that remained significantly downregulated in both CPB and recovery, and cluster 3 (purple) represents the miRNAs that were significantly downregulated in at least one timepoint during CPB and significantly upregulated in at least one timepoint in recovery. We used the TargetScan database to identify putative mRNA targets of the miRNAs within each of the three clusters. The number of genes associated with each miRNA and within each cluster is displayed in **Supplemental Table S2**. We filtered out genes that were not present in a parallel mRNA sequencing analysis done by our group^4^. (Figure 1C).

In a previous study using a separate population, we identified 2, 688 genes which exhibited temporal dysregulation after CPB, which were categorized into 5 distinct groups.^4^ We expanded on this analysis to investigate relationships between these differentially expressed genes (DEGs) and the miRNA target genes identified in this analysis. 1,046 of these 2,688 DEGs (38.9%) were reported targets of differentially expressed miRNAs, as shown in **Table 2 and Figure 1C**. 70 DEGs were targeted by miRNAs in all three clusters, 48 DEGs were targeted by miRNAs in clusters 1 and 3, 60 mRNAs were targeted by miRNAs in clusters 1 and 2, 208 DEGs were targeted by miRNAs in clusters 2 and 3, 75 DEGs were targeted by miRNAs in cluster 1 alone, 250 DEGs were targeted by miRNAs in cluster 3 alone, and 335 DEGs were targeted by miRNAs in cluster 2 alone (**see Figure 1D**). Many of these DEGs were targeted by multiple miRNAs within the same cluster. miRNA target genes contained DEGs across all five mRNA categories (Table 2), with most miRNAs across all clusters targeting genes in the “blue” category (indicating gene expression downregulated across all timepoints), or “red” category (i.e., gene expression increased then decreased). Thus, these changes in miRNA expression may be attributable to some of the previously observed changes in gene expression after CPB.

**Table 2 –.**
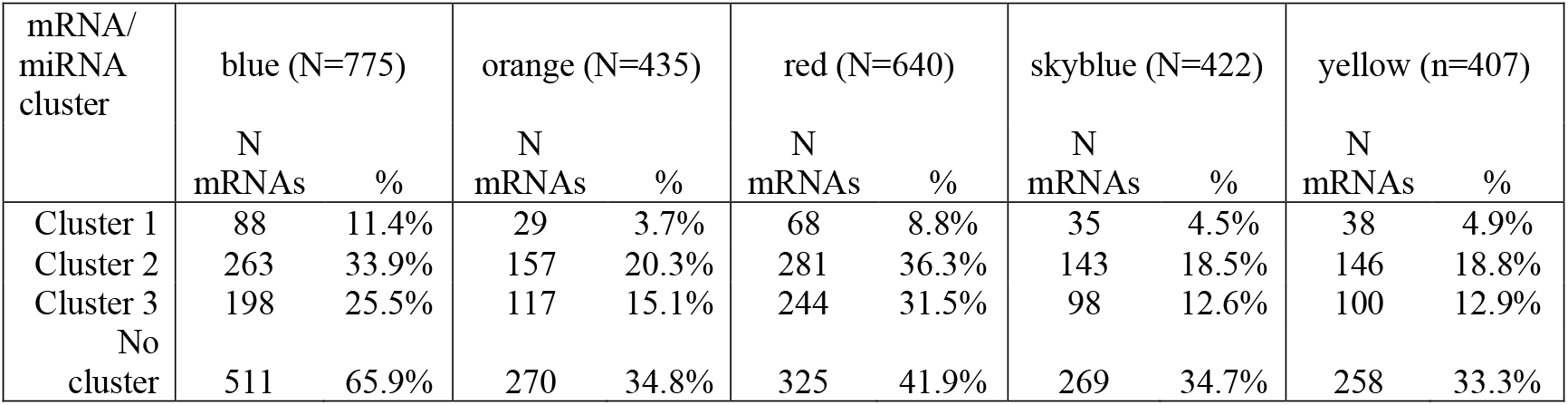
Proportion of target genes in each miRNA cluster which were differentially expressed in Previous Study. ^4^.

| mRNA/<br>miRNA<br>cluster | blue (N=775) |  | orange (N=435) |  | red (N=640) |  | skyblue (N=422) |  | yellow (n=407) |  |
| --- | --- | --- | --- | --- | --- | --- | --- | --- | --- | --- |
|  | N<br>mRNAs | % | N<br>mRNAs | % | N<br>mRNAs | % | N<br>mRNAs | % | N<br>mRNAs | % |
| Cluster 1 | 88 | 11.4% | 29 | 3.7% | 68 | 8.8% | 35 | 4.5% | 38 | 4.9% |
| Cluster 2 | 263 | 33.9% | 157 | 20.3% | 281 | 36.3% | 143 | 18.5% | 146 | 18.8% |
| Cluster 3 | 198 | 25.5% | 117 | 15.1% | 244 | 31.5% | 98 | 12.6% | 100 | 12.9% |
| No<br>cluster | 511 | 65.9% | 270 | 34.8% | 325 | 41.9% | 269 | 34.7% | 258 | 33.3% |

### KEGG pathway and target gene analysis

In order to identify biologic pathways that were modulated by miRNA clusters, we performed KEGG biological pathways analysis. For this analysis, we used the top 200 target genes for every miRNA based on context score. Overall, we observed that the target genes of these miRNAs were involved in the cell growth and death, cellular community, immune, and signal transduction KEGG subgroups (Figure 2). MiRNAs from cluster 1 targeted 512 genes that were enriched with the p53 signaling pathway and the cell cycle pathway (FDR Adjusted p <0.05), which are both involved in cellular growth and death (**Figure 2**). MiRNAs from cluster 2 targeted 1708 genes that were enriched within 32 different biological pathways (**Figure 2**). Several pathways related to immunity—such as phagocytosis, B-cell receptors, and T-cell receptors, were identified as being modulated by this cluster The most significantly enriched pathway was for neurotrophin signaling. Nine of these pathways were signal transduction pathways, and six of these pathways were pathways related to the immune system (**Figure 2**). MiRNAs from cluster 3 targeted 1430 genes that were enriched with 5 different biological pathways. We also examined similarities and differences in the pathways targeted by these miRNAs. The p53 signaling pathway was enriched for genes which were targets of miRNAs in both cluster 1 and cluster 2. There were no overlapping pathways between clusters 1 and 3. There were 4 pathways which overlapped between clusters 2 and 3. Overall The most statistically significant pathway for each cluster were those related to the P53 signaling pathway (FDR adjusted *p* =4.77 x 10^-5^, Cluster l), the Fc gamma R-mediated phagocytosis pathway (FDR adjusted 3.44×10^-4^, Cluster 2) (**Figure 2**), and the neurotrophin signaling pathway (FDR adjusted 6.84 x 10^-3^, Cluster 3). From our previous research, we have demonstrated that MEK/ERK signaling is important for CPB induced expression of IL8 and TNFa ^4^. miRNAs from cluster 2 were predicted targets of 38 genes in the MAPK pathway, and this pathway was significantly enriched for these genes (p<0.05). Some of the miRNA target genes included regulatory kinases of differentially expressed mRNAs found in our corresponding mRNA-seq data (**Figure 3**).^4^

**Figure 2.**
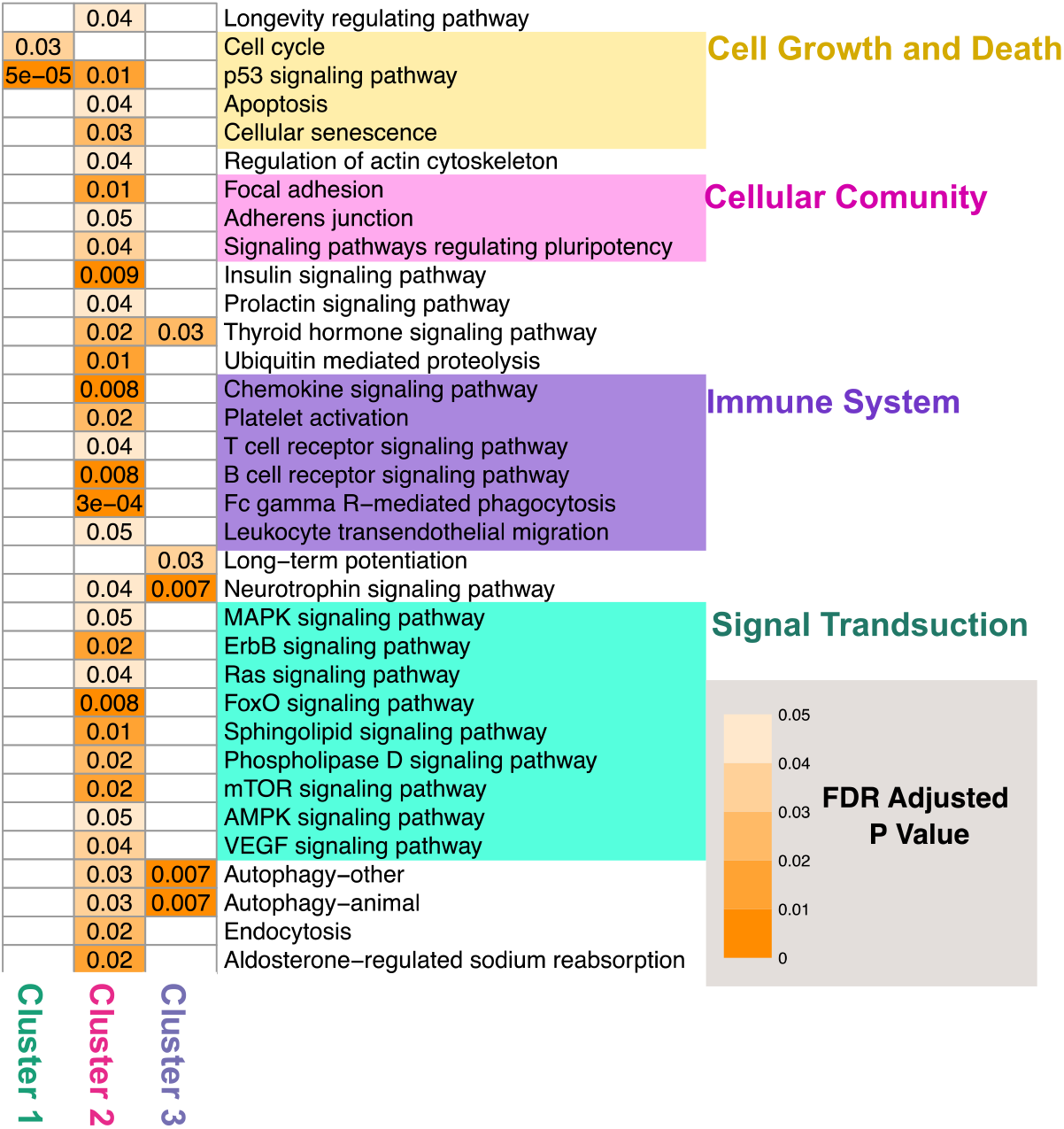
Pathway enrichment analysis of the miRNA target genes in each cluster. The heatmap showing significant KEGG biological pathways enriched for the top genes regulated by the miRNAs in each cluster. The numbers indicate the adjusted p-values for the pathways.

**Figure 3.**
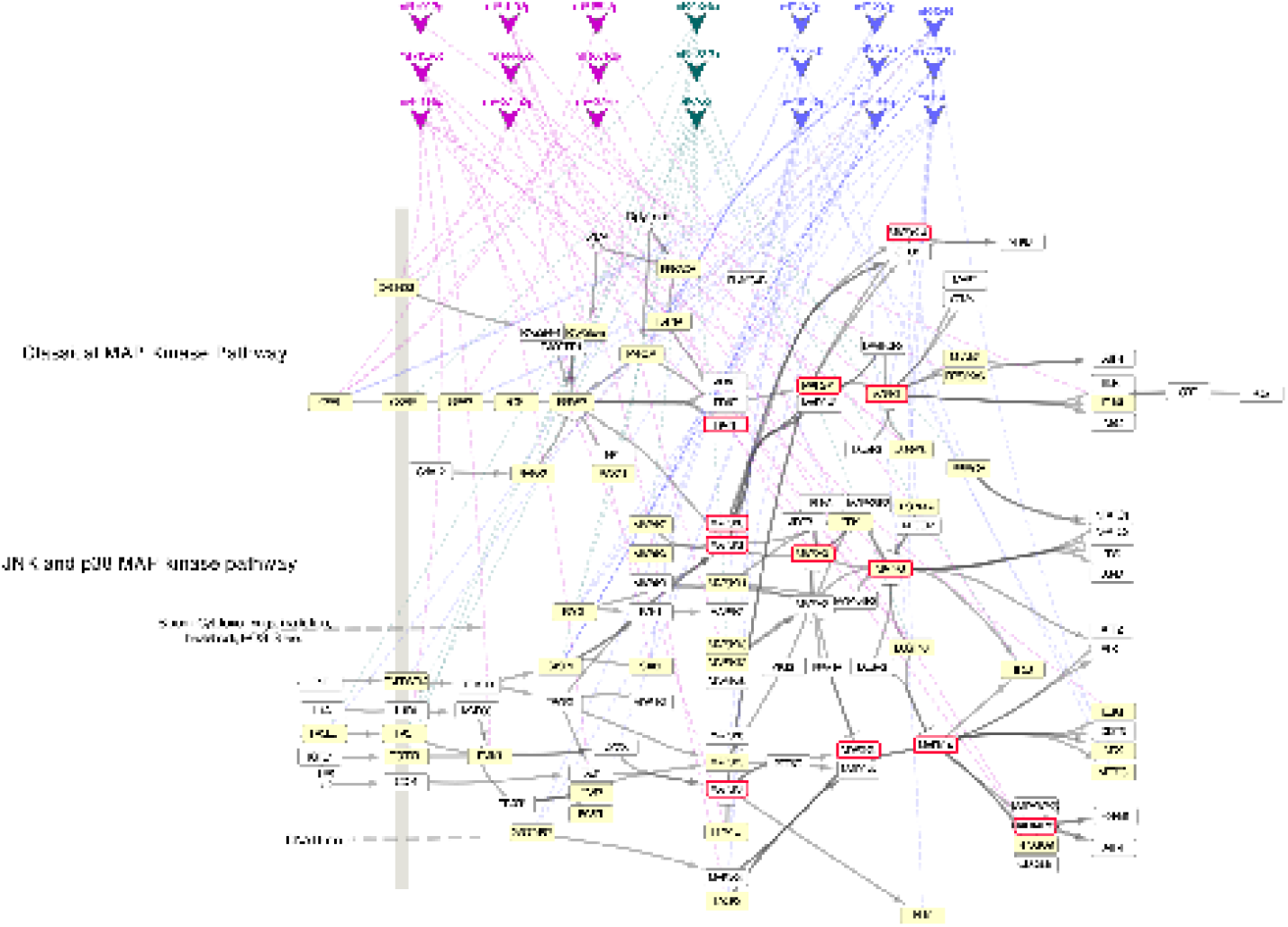
Targeted genes in the Mitogen-Activated Protein Kinase pathway. Genes in the MAPK pathway which are targets of the differentially expressed miRNAs are highlighted in yellow. Genes predicted as the regulatory kinases of differentially expressed mRNAs in the matched mRNA-seq dataset are highlighted with the red border.

### Cytokine-Receptor Network Analysis

Since the many KEGG pathways enriched for the miRNA target genes involved immune signaling, including the chemokine signaling pathway, we generated a subnetwork of all receptor-ligand network reactions for our miRNA target genes using the FANTOM5 database.^17^ Overall, cluster 1 was a potential regulator of 17 ligands and 11 receptors; cluster 3 was a potential regulator of 43 ligands and 31 receptors, and cluster 2 was a potential regulator of 47 receptors and 25 ligands (see **Supplemental Table S2**). We performed a targeted analysis by looking at different mediators of inflammation associated with each of the miRNA targets from each cluster. We filtered target genes of each miRNA by cytokines and their receptors (**Figure 4A**), chemokines and their receptors (**Fig 4B**), and integrin receptors and adhesion molecules (**Figure 4C**). The color code of each miRNA denotes which cluster each is derived from.

**Figure 4.**
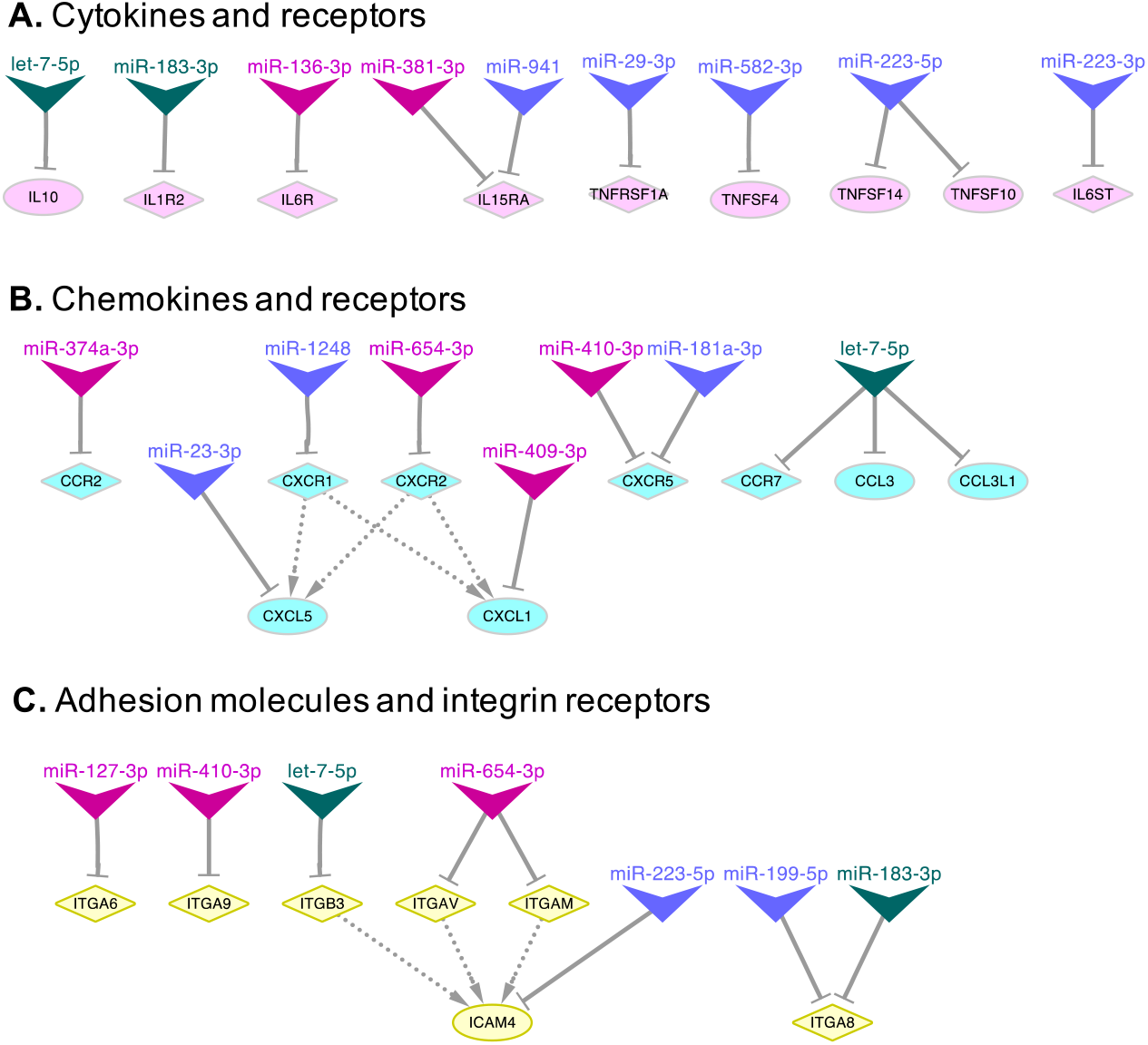
Receptor-ligand networks for the target genes in each cluster. Analysis of receptor-ligand networks were performed for target genes using the Fantom5 database. The networks of cytokines and receptors **(A)**, chemokines and receptors **(B)** and integrin receptors and adhesion molecules **(C)** targeted by the miRNAs in each cluster were demonstrated.

## DISCUSSION

miRNAs play important roles in post-transcriptional regulation of target genes. Thus, understanding how miRNA levels in leukocytes fluctuate during and after CPB significantly may help to both advance the understanding of the mechanisms that underpin CPB associated inflammation, and develop biomarkers of CPB related complications. To our knowledge, this study is the first to characterize the novel global miRNA changes that occur in circulating leukocytes of CPB patients. The dataset presented here significantly advanced the state of knowledge since the previously reported studies (**Table S3** have examined only a limited number of miRNAs change in CPB patients using mainly serum or plasma samples. Our study specifically examined changes in circulating nucleated cells, a cell population that is an important driver of inflammation. In addition, the global profiling of all miRNA changes in an unbiased manner significantly advances the understanding of how circulating leukocytes are responding to CPB. The data presented in this study, especially regarding the miRNAs that are predicted to facilitate the resolution of inflammation, may form the basis of novel therapeutic and risk stratification strategies.

Given that miRNA typically reduce the protein levels of their target genes^6^, our data suggest that miRNAs are involved in the development and resolution of CPB associated inflammation. During CPB, the changes in miRNA levels promote inflammation. miRNAs that are upregulated during CPB (Cluster 1) target genes that typically are anti-inflammatory, such as IL-10 and IL1R2 (Figure 4A) At the same time miRNAs that are down regulated during CPB (Clusters 2 and 3) target receptors for key inflammatory signaling pathways, such as TNF-α and IL-8 (**Figure 4A** and **B**). These changes have the net result of increasing the activity of inflammatory pathways.

The role of the members of Cluster 2, miRNAs that are decreased during and after CPB is especially interesting given this cluster is predicted to uniquely modulate pathways and receptors involved with inflammation. As these miRNAs generally are thought to decrease gene expression, reduction of expression of these negative regulators during and after surgery may lead to corresponding increased expression of their target genes involved in immune regulation. As shown in **Figure 2**, members of this cluster are predicted to repress pathways such as leukocyte migration, chemokine signaling, and MAPK signaling. Cytokines that are activated during CPB^4, 19–25^—such as IL6, IL8, and TNF-α, or their receptors are targeted by members of this cluster of miRNAs.

Cluster 3, miRNAs that are decreased during CPB and rise after CPB, modulate genes related to cytokine signaling. For example, miR-223 (A member of Cluster 3) has been shown to target interleukin-6 (IL6)^26^. miR-23a is a miRNA that targets and inhibits pro-inflammatory cytokine interleukin-8 (IL8) via signal transducer and activator of transcription 3 (STAT3) inhibition^27^. The IL-8 receptor, CXCR1, is predicted to be targeted by miR-1248 while another of its receptors, *CXCR2* is targeted by miR-223^26^. miRNA-29-3p ^28^ and miR-223-5p^29^ target *TNFRSF1*, which encodes a key receptor for TNF signaling. Based upon these data, we believe that the members of miRNA cluster 3 play important roles in modulating CPB-associated inflammation in circulating leukocytes.

Our study is limited in the small sample size of our patient samples; however, there are clear patterns of specific miRNA differential expression amongst all patient samples, based on dense temporal mapping and comparisons within each patient. We also base our analyses on the assumption that miRNAs negatively regulate gene expression, when in reality this relationship is more complex.^6^ More research is necessary to validate and isolate specific functions of these miRNAs. Some of the strengths of this study are the global miRNA profiling of circulating miRNAs at several timepoints during and after CPB. It is important to expand our understanding of miRNA role in the context of CPB-induced inflammation to help us further uncover all the mechanisms in which inflammation is induced and operates. Furthermore, miRNAs may also have the potential to be a non-invasive, non-pharmacologic treatment in moderating this inflammation.

## WORKS CITED

1. Kansy A, Tobota Z, Maruszewski P, Maruszewski B. Analysis of 14,843 neonatal congenital heart surgical procedures in the European Association for Cardiothoracic Surgery Congenital Database. Ann Thorac Surg. 2010;89:1255–1259.

2. Smith AH, Gay JC, Patel NR. Trends in resource utilization associated with the inpatient treatment of neonatal congenital heart disease. Congenit Heart Dis. 2014;9:96–105.

3. Appachi E, Mossad E, Mee RB, Bokesch P. Perioperative serum interleukins in neonates with hypoplastic left-heart syndrome and transposition of the great arteries. J Cardiothorac Vasc Anesth. 2007;21:184–190.

4. Tu LN, Hsieh L, Kajimoto M, et al. Shear stress associated with cardiopulmonary bypass induces expression of inflammatory cytokines and necroptosis in monocytes. JCI insight. 2021;6.

5. Kehl T, Backes C, Kern F, et al. About miRNAs, miRNA seeds, target genes and target pathways. Oncotarget. 2017;8:107167–107175.

6. O’Brien J, Hayder H, Zayed Y, Peng C. Overview of MicroRNA Biogenesis, Mechanisms of Actions, and Circulation. Front Endocrinol (Lausanne). 2018;9:402.

7. Momen-Heravi F, Bala S. miRNA regulation of innate immunity. J Leukoc Biol. 2018.

8. Kiss A, Heber S, Kramer AM, et al. MicroRNA Expression Profile Changes after Cardiopulmonary Bypass and Ischemia/Reperfusion-Injury in a Porcine Model of Cardioplegic Arrest. Diagnostics (Basel). 2020;10.

9. Abu-Halima M, Poryo M, Ludwig N, et al. Differential expression of microRNAs following cardiopulmonary bypass in children with congenital heart diseases. J Transl Med. 2017;15:117.

10. Allen RM, Zhao S, Ramirez Solano MA, et al. Bioinformatic analysis of endogenous and exogenous small RNAs on lipoproteins. J Extracell Vesicles. 2018;7:1506198.

11. Kechin A, Boyarskikh U, Kel A, Filipenko M. cutPrimers: A New Tool for Accurate Cutting of Primers from Reads of Targeted Next Generation Sequencing. J Comput Biol. 2017;24:1138–1143.

12. Langmead B, Trapnell C, Pop M, Salzberg SL. Ultrafast and memory-efficient alignment of short DNA sequences to the human genome. Genome Biol. 2009;10:R25.

13. Love MI, Huber W, Anders S. Moderated estimation of fold change and dispersion for RNA-seq data with DESeq2. Genome Biol. 2014;15:550.

14. Benjamini Y, Hochberg Y. Controlling the False Discovery Rate: A Practical and Powerful Approach to Multiple Testing. Journal of the Royal Statistical Society: Series B (Methodological). 1995;57:289–300.

15. Agarwal V, Bell GW, Nam JW, Bartel DP. Predicting effective microRNA target sites in mammalian mRNAs. Elife. 2015;4.

16. Kuleshov MV, Jones MR, Rouillard AD, et al. Enrichr: a comprehensive gene set enrichment analysis web server 2016 update. Nucleic Acids Res. 2016;44:W90–97.

17. Shannon P, Markiel A, Ozier O, et al. Cytoscape: a software environment for integrated models of biomolecular interaction networks. Genome Res. 2003;13:2498–2504.

18. Ramilowski JA, Goldberg T, Harshbarger J, et al. A draft network of ligand-receptor-mediated multicellular signalling in human. Nat Commun. 2015;6:7866.

19. Butler J, Pathi VL, Paton RD, et al. Acute-phase responses to cardiopulmonary bypass in children weighing less than 10 kilograms. Ann Thorac Surg. 1996;62:538–542.

20. Ashraf SS, Tian Y, Cowan D, et al. Proinflammatory cytokine release during pediatric cardiopulmonary bypass: influence of centrifugal and roller pumps. J Cardiothorac Vasc Anesth. 1997;11:718–722.

21. Hauser GJ, Ben-Ari J, Colvin MP, et al. Interleukin-6 levels in serum and lung lavage fluid of children undergoing open heart surgery correlate with postoperative morbidity. Intensive Care Med. 1998;24:481–486.

22. Bokesch PM, Kapural MB, Mossad EB, et al. Do peritoneal catheters remove pro-inflammatory cytokines after cardiopulmonary bypass in neonates? Ann Thorac Surg. 2000;70:639–643.

23. Hovels-Gurich HH, Vazquez-Jimenez JF, Silvestri A, et al. Production of proinflammatory cytokines and myocardial dysfunction after arterial switch operation in neonates with transposition of the great arteries. J Thorac Cardiovasc Surg. 2002;124:811–820.

24. Alcaraz AJ, Manzano L, Sancho L, et al. Different proinflammatory cytokine serum pattern in neonate patients undergoing open heart surgery. Relevance of IL-8. J Clin Immunol. 2005;25:238–245.

25. Gu CH, Cui Q, Wang YY, et al. Effects of insulin therapy on inflammatory mediators in infants undergoing cardiac surgery with cardiopulmonary bypass. Cytokine. 2008;44:96–100.

26. Dorhoi A, Iannaccone M, Farinacci M, et al. MicroRNA-223 controls susceptibility to tuberculosis by regulating lung neutrophil recruitment. J Clin Invest. 2013;123:4836–4848.

27. Qu JQ, Yi HM, Ye X, et al. MiR-23a sensitizes nasopharyngeal carcinoma to irradiation by targeting IL-8/Stat3 pathway. Oncotarget. 2015;6:28341–28356.

28. Gao XZ, Zhang ZX, Han GL. MiR-29a-3p Enhances the Viability of Rat Neuronal Cells that Injured by Oxygen-Glucose Deprivation/Reoxygenation Treatment Through Targeting TNFRSF1A and Regulating NF-kappaB Signaling Pathway. J Stroke Cerebrovasc Dis. 2020;29:105210.

29. Qin D, Wang X, Li Y, et al. MicroRNA-223-5p and −3p Cooperatively Suppress Necroptosis in Ischemic/Reperfused Hearts. J Biol Chem. 2016;291:20247–20259.

30. Bolkier Y, Nevo-Caspi Y, Salem Y, Vardi A, Mishali D, Paret G. Micro-RNA-208a, −208b, and-499 as Biomarkers for Myocardial Damage After Cardiac Surgery in Children. Pediatr Crit Care Med. 2016;17:e193–197.

31. Zloto K, Tirosh-Wagner T, Bolkier Y, et al. MiRNA-208a as a Sensitive Early Biomarker for the Postoperative Course Following Congenital Heart Defect Surgery. Pediatric cardiology. 2018;39:1565–1571.

32. Poon KS, Palanisamy K, Chang SS, et al. Plasma exosomal miR-223 expression regulates inflammatory responses during cardiac surgery with cardiopulmonary bypass. Sci Rep. 2017;7:10807.

33. Kang Z, Li Z, Huang P, et al. Remote ischemic preconditioning upregulates microRNA-21 to protect the kidney in children with congenital heart disease undergoing cardiopulmonary bypass. Pediatr Nephrol. 2018;33:911–919.

34. Zhou X, Mao A, Wang X, Duan X, Yao Y, Zhang C. Urine and serum microRNA-1 as novel biomarkers for myocardial injury in open-heart surgeries with cardiopulmonary bypass. PLoS One. 2013;8:e62245.

35. Mazzone AL, Baker RA, McNicholas K, Woodman RJ, Michael MZ, Gleadle JM. Circulating and Urinary miR-210 and miR-16 Increase during Cardiac Surgery Using Cardiopulmonary Bypass - A Pilot Study. The journal of extra-corporeal technology. 2018;50:19–29.

36. Yang K, Gao B, Wei W, et al. Changed profile of microRNAs in acute lung injury induced by cardio-pulmonary bypass and its mechanism involved with SIRT1. International journal of clinical and experimental pathology. 2015;8:1104–1115.

37. Yao Y, Du J, Cao X, et al. Plasma levels of microRNA-499 provide an early indication of perioperative myocardial infarction in coronary artery bypass graft patients. PloS one. 2014;9:e104618.

