## Supplemental Material for "microRNA Expression Levels Change in Neonatal Patients During and After Exposure to Cardiopulmonary Bypass"

**SUPPLEMENTAL FIGURES**

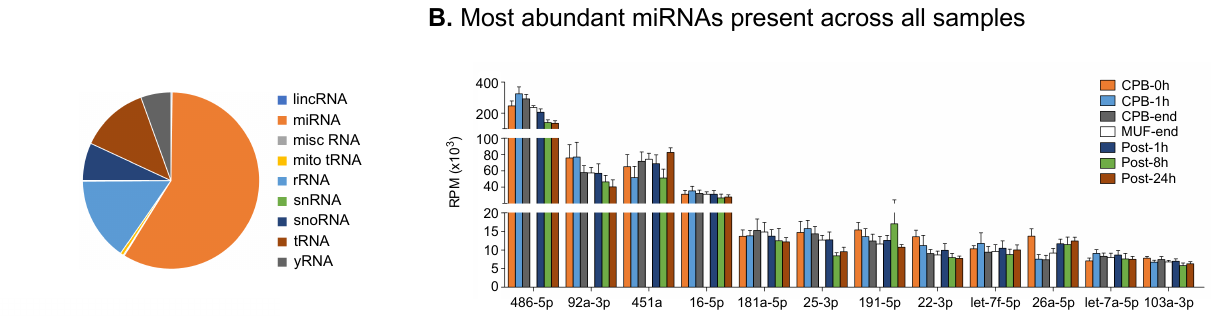

**SUPPLEMENTAL TABLES**

**Table S1.The number of miRNA gene targets used in sub-analyses.**

| **miRNA** | **Cluster** | **Target Genes (Targetscan)** | **Target Genes with \|Context Score\|>0** | **Target Genes Present in mRNA Analysis*** | **Target Genes significantly DE*** | **Genes used for KEGG Pathway Enrichment** |
| --- | --- | --- | --- | --- | --- | --- |
| miR-183-3p | C1 | 2599 | 1722 | 1128 | 194 | 200 |
| let-7-5p |  | 332 | 332 | 237 | 41 | 200 |
| miR-182-5p |  | 172 | 172 | 119 | 23 | 119 |
| miR-654-3p | C2 | 2523 | 1921 | 1264 | 232 | 200 |
| miR-27-3p |  | 1092 | 1055 | 751 | 135 | 200 |
| miR-136-3p |  | 1155 | 1137 | 726 | 128 | 200 |
| miR-374a-3p |  | 1355 | 737 | 487 | 73 | 200 |
| miR-144-5p |  | 938 | 715 | 485 | 79 | 200 |
| miR-410-3p |  | 604 | 558 | 395 | 68 | 200 |
| miR-409-3p |  | 316 | 286 | 199 | 21 | 199 |
| miR-381-3p |  | 128 | 128 | 93 | 13 | 93 |
| miR-411-5p.1 |  | 123 | 118 | 89 | 7 | 89 |
| miR-127-3p |  | 25 | 25 | 17 | 1 | 17 |
| miR-1248 | C3 | 2843 | 2013 | 1347 | 223 | 200 |
| miR-582-3p |  | 1622 | 1622 | 1101 | 174 | 200 |
| miR-29-3p |  | 755 | 755 | 562 | 112 | 200 |
| miR-23-3p |  | 713 | 713 | 517 | 87 | 200 |
| miR-199-5p |  | 634 | 610 | 462 | 64 | 200 |
| miR-223-5p |  | 994 | 662 | 432 | 83 | 200 |
| miR-223-3p |  | 415 | 405 | 329 | 60 | 200 |
| miR-941 |  | 387 | 362 | 222 | 41 | 200 |
| miR-181a-3p |  | 322 | 322 | 201 | 49 | 200 |
| miR-24-3p |  | 119 | 119 | 81 | 14 | 81 |

**Table S2 – Ligands and receptors as target genes of differentially expressed miRNAs in each cluster**

| **Cluster** | **miRNA** | **Receptors** | **Ligands** |
| --- | --- | --- | --- |
| Cluster 1 | let-7-5p | *TGFBR1, ADRB2, FAS, CCR7, IGF1R, PGRMC1, ITGB3, TGFBR3, FAS* | *EDN1, IL10, S100A8, CCL3L1, FASLG, CCL3, CCL3L3* |
|  | miR-182-5p | *EDNRB, VLDLR* | *HBEGF* |
|  | miR-183-3p | *IL1R2, LEPR, IFNGR1, ASGR2, KLRD1, ITGA8, CD109* | *BMP3, LPL, LYZ* |
| Cluster 3 | miR-199-5p | *DDR1, BCAM, CELSR1, FZD6, ITGA8, ACVR1B, PLXNC1, ACVR1B, ACVR1B* | *LIN7C, HG* |
|  | miR-582-3p | *TFRC, BMPR2, TFRC* | *TNFSF4* |
|  | miR-223-3p | *IL6ST, ACVR2A, SCARB1, SDC2* | *HSP90B1, SORBS1* |
|  | miR-223-5p | *DPP4, CD177, ERAP1, LRP8, DPP4, CD177, ERAP1, LRP8* | *TNFSF14, TIMP3, TNFSF10, OLAH, ICAM4, HSPA1A, EFNB2, FBN1* |
|  | miR-29-3p | *ROBO1, TNFRSF1A, CNR1, F11R, CNR1* | *COL1A1, COL3A1, COL7A1, PDGFC, VEGFA, COL6A3* |
|  | miR-23-3p | *ABCA1, NRXN3* | *PROK2, CXCL5* |
|  | miR-181a-3p | *CD93, EDNRB, CXCR5, VANGL1, F2R* | *COL9A2, APP, ADAM10* |
|  | miR-24-3p | *LMBR1L, SIRPA* | *FASLG, LAMB3* |
|  | miR-1248 | *KLRK1, CD81, EPHB6, CD28, CXCR1, ADCY7, LEPR* | *HBEGF, HMGB1, PDGFB, DUSP18* |
|  | miR-941 | *ORAI2, IFNAR1, GFRA2, MGRN1, TLR6, IL15RA* | *TIMP2* |
| Cluster 2 | miR-127-3p | *ITGA6* | *DLK1* |
|  | miR-136-3p | *ERAP1, IL6R, NPTN, CANX, NPTN* | *CLCF1* |
|  | miR-144-5p | *TMEM67, F2R, TLR2, TFR2, CD3G* | *RELN* |
|  | miR-27-3p | *ADORA2B, NRP2, CD28, ST14, BMPR2* | *HBEGF, COLQ, SEMA6A* |
|  | miR-381-3p | *IL15RA, LRRC4, SLC18A2* | *PDAP1, PDGFC* |
|  | miR-409-3p | *S1PR1, LRP6, PLXNC1, TGFBR3, MYLK, ACVR2B* | *CXCL1, CALM1, LIN7C, SEMA4F* |
|  | miR-410-3p | *NRXN3, PLXNA2, ITGA9, CXCR5, MCAM, IFNAR2* | *HMGB1, ADM, RGMB* |
|  | miR-411-5p | *TGFBR2* | *CLEC11A* |
|  | miR-654-3p | *CXCR2, TFRC, DYSF, OLR1, ITGAM, ITGAV, SLC37A1, KCNJ15, KLRC1, CNR1, KLRC2, PLXNA4* | *MST1, FLT3LG, APP* |
|  | miR-374a-3p | *KLRG1, LGR4, AMFR, CCR2* | *VEGFB, CALM3, LTBP3, EDIL3, B2M, LPL* |

**Table S3 – Plasma and serum miRNAs reported in the literature as biomarkers for complications in CPB surgery**

| **miRNA** | **Age** | **Elevation time point** | **Associated complications** | **Reference** |
| --- | --- | --- | --- | --- |
| miR-208a/b  miR-499 | Pediatric | 6-24h post CPB | Myocardial injury  Length of hospital stay | ^30, 31^ |
| miR-223 | Adult | 2-6h after start of CPB | Inflammation | ^32^ |
| miR-21 | Pediatric | 6-24h post CPB | Acute kidney injury | ^33^ |
| miR-1 | Adult | 1-24h post CPB | Myocardial injury | ^34^ |
| miR-210  miR-16 | Adult | CPB & 4h post CPB | Myocardial injury | ^35^ |
| miR-320  miR-200c  miR-205 | Adult | 8-16h post CPB | Acute lung injury | ^36^ |
| miR-133a  miR-499 | Adult | 1-6h post CPB | Myocardial injury | ^37^ |
